# Accelerated multi-shell diffusion MRI with Gaussian process estimated reconstruction of multi-band imaging

**DOI:** 10.1101/2024.10.08.617197

**Authors:** Xinyu Ye, Karla Miller, Wenchuan Wu

## Abstract

**Purpose:** This work aims to propose a robust reconstruction method exploiting shared information across shells to increase the acquisition speed of multi-shell diffusion MRI, enabling rapid tissue microstructure mapping.

**Theory and Methods:** Local q-space points share similar information. Gaussian Process can exploit the q-space smoothness in a data-driven way and provide q-space signal estimation based on the signals from a q-space neighborhood. The Diffusion Acceleration with Gaussian process Estimated Reconstruction (DAGER) method uses the signal estimation from Gaussian process as a prior in a joint k-q reconstruction and improves image quality under high acceleration factors compared to conventional (k-only) reconstruction. In this work, we extend the DAGER method by introducing a multi-shell covariance function and correcting for Rician noise distribution in magnitude data when fitting the Gaussian process model. The method was evaluated with both simulation and in vivo data.

**Results:** Simulated and in-vivo results demonstrate that the proposed method can significantly improve the image quality of reconstructed dMRI data with high acceleration both in-plane and slice-wise, achieving a total acceleration factor of 12. The improvement of image quality allows more robust diffusion model fitting compared to conventional reconstruction methods, enabling advanced multi-shell diffusion analysis within much shorter scan time.

**Conclusion:** The proposed method enables highly accelerated dMRI which can shorten the scan time of multi-shell dMRI without sacrificing quality compared to conventional practice. This may facilitate a wider application of advanced dMRI models in basic and clinical neuroscience.

## Introduction

Diffusion-weighted MRI (dMRI) is a non-invasive technique that can characterize tissue microstructure and delineate structural connectivity by measuring the movement of water within biologically tissues, making it an essential tool for basic and clinical neuroscience research [1-5]. The acquisition time of a dMRI scan is determined by the number of diffusion directions (i.e., q-space samples) and the scan time per direction (acquiring k-space). Due to the desirability of both high spatial and angular resolution (k- and q-space, respectively), image acceleration has become critical to dMRI acquisition. Advanced diffusion MRI models (e.g., DKI, NODDI, CHARMED [6-8]) provide higher specificity about tissue microstructure compared to simple models (e.g., DTI). However, fitting these advanced models requires acquisitions of a large number of q-space samples over many diffusion directions and across multiple b shells, leading to long scan times [9-11]. dMRI signal is typically acquired with 2D single-shot EPI due to its rapid acquisition speed, for which the scan time of each q-space sample is coupled with the number of slices. Consequently, achieving higher resolution (thinner slices) with the same brain coverage results in longer scan time per direction. When the total scan time is limited, this creates a fundamental tradeoff between spatial resolution, anatomical coverage, and model specificity.

Various acquisition and reconstruction methods have been developed to speed up dMRI scans and mitigate this tradeoff. Conventional parallel imaging methods such as SENSE and GRAPPA [12-13] have been applied to accelerate dMRI by under-sampling in k space. However, as more than 50% of dMRI sequence timing is dedicated to diffusion sensitizing gradients rather than readout, in-plane under-sampling is not as effective in reducing scan time. A major advance in dMRI acceleration is Simultaneous multislice (SMS) imaging [14], which reduces the scan time by acquiring multiple slices at the same time and thus effectively sharing the diffusion sensitizing gradients across slices. With blipped-CAIPI sampling [15] SMS allows 2-3X slice acceleration for dMRI without significantly increased g-factor SNR penalties. However, due to the intrinsically low SNR of dMRI data, achieving high slice acceleration (i.e., 4 or higher) is still challenging. This is particularly problematic when combined with in-plane accelerations because the two approaches compete for the use of coil sensitivity information, leading to a highly ill-conditioned reconstruction problem. Methods enforcing some prior information in the spatial domain can potentially be employed to alleviate the problem [16-18]. But the slice acceleration achievable in dMRI is still highly limited compared to other MRI modalities. Using large cohort studies as an example, the UK Biobank [19] and Human Connectome Project [20] use a slice acceleration factor of 2 or 3 for dMRI, but an acceleration factor of up to 8 for functional MRI.

Acceleration methods have also been developed to reduce the number of required q-space directions. To exploit the data redundancy, different methods like compressed sensing [21-22] and dictionary learning [23] have been used to improve the estimation of diffusion metrics from a reduced number of q-space samples. Deep learning-based methods have been also developed to improve the accuracy of diffusion metrics estimation from a reduced number of diffusion directions [24-27].

Finally, joint k-q acceleration methods have been proposed to leverage the information redundancy in both k-space and q-space simultaneously to achieve a higher acceleration factor [28-35] by jointly reconstructing k-space data at different q space locations. Diffusion Acceleration with Gaussian process Estimated Reconstruction (DAGER) [35] is a model-independent joint k-q reconstruction method that exploits the smoothness in q-space to interpolate signals in a data-driven way, which enables high acceleration factor. DAGER uses Gaussian-Processes (GPs) to model dMRI signals, an approach that has been successfully applied to correct for subject motion and eddy current distortions [36]. Whereas some other joint k-q reconstruction methods use priors in the context of a specific signal model, DAGER does not make assumptions about the form of dMRI signal. It also doesn’t impose spatial-smoothing priors in the reconstruction but only data-driven q-space smoothness fitted by Gaussian Processes. Besides, DAGER uses a Bayesian framework to automatically estimate the reconstruction hyperparameters from the data. This avoids the manual tuning of regularization parameters.

However, the original DAGER method only exploits information redundancy between different diffusion directions within a single shell in q-space (i.e., a single b-value). For multi-shell dMRI acquisitions, DAGER currently reconstructs each shell independently, which may require a large number of diffusion directions. Since signal also varies smoothly with q-space radius in addition to angle, exploiting information across shells in the reconstruction can reduce the needed number of diffusion directions for each shell and improve the image quality particularly for outer q-shells due to their lower SNR. In this work, we propose to extend the GP model used in the original DAGER to enable information sharing between shells to improve reconstruction at high acceleration for multi-shell acquisition. In addition, the original DAGER framework assumes Gaussian noise, which may underestimate the noise variance given that magnitude data (with a Rician distribution) is used in GP fitting. This assumption is particularly problematic for low-SNR data, such as at high b-values. We integrate a variance-stabilizing transformation [37-38] to improve the accuracy of Gaussian Process fitting under Rician noise conditions. A data-driven multi-shell diffusion reconstruction method is developed that integrates these two extensions of DAGER. The method, which we name multi-shell DAGER (ms-DAGER), is evaluated using numerical simulations and in vivo scans. We demonstrate robust image quality and microstructure model fitting at 1.25-1.5mm resolution using high multi-slice and in-plane accelerations (SMS=4 × R=3).

## Theory

### GP modelling of dMRI signal

A Gaussian process (GP) is a data-driven model that, rather than fitting a single function, produces an infinite set of functions that all describe the data. Specifically, each function fits the data exactly at the measured points and the collective set of functions follow a multi-variate Gaussian distribution away from the measured points [39]. dMRI signals exhibit smoothness in q-space as shown in Figure 1.(a) – that is, they have a covariance structure – that can be described using a multi-variate Gaussian. GP models are able to capture this structure in a way that enables the prediction of the mean and variance of the dMRI signal at unseen q-space locations. With GP modelling, we assume that dMRI signals ***u*** at a set of 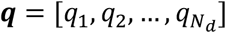 follows a multivariate Gaussian distribution:

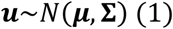

which is determined by a mean vector μ (*N*_d_ x 1) and a covariance matrix **Σ** (*N*_d_ x *N*_d_). Here, ***u*** represents dMRI signals from a single voxel for all dMRI directions and *N*_d_ is the number of dMRI directions.

**Fig 1.**
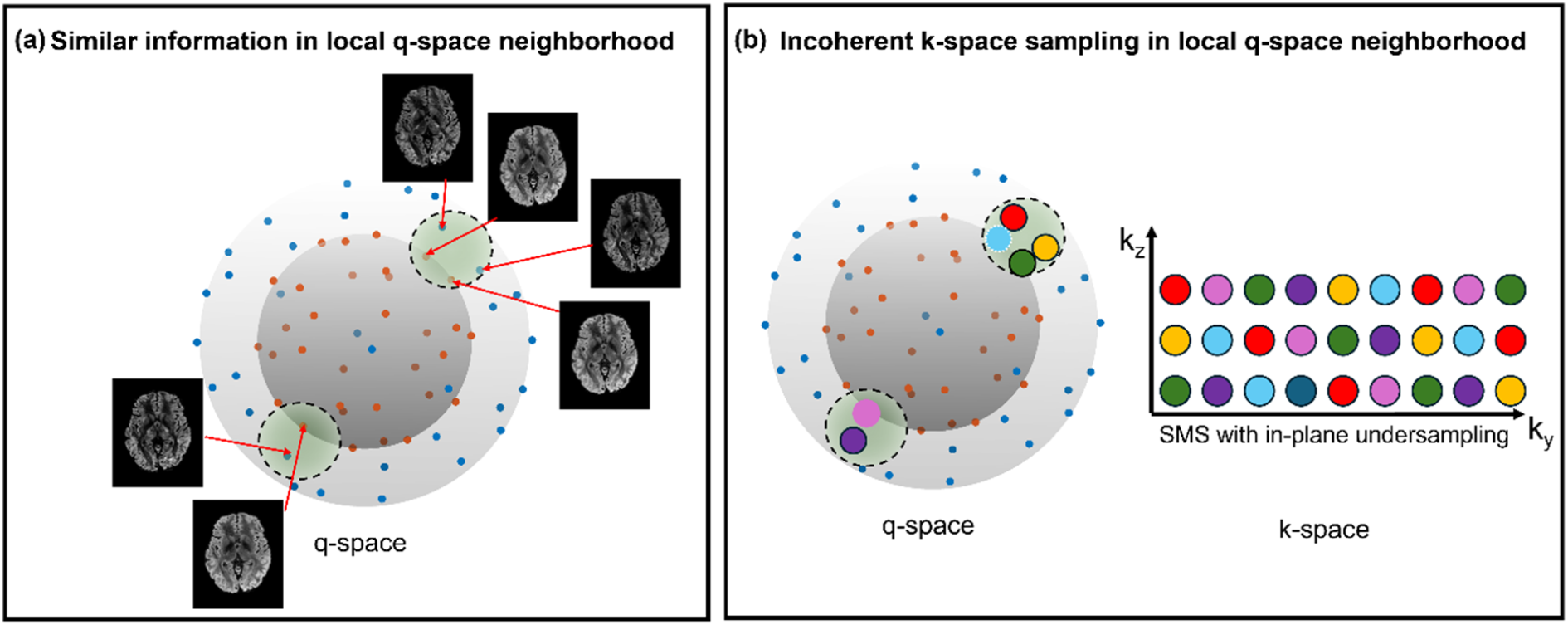
ms-DAGER reconstruction utilizes local smoothness in q-space. a) dMRI images that are near to each other in q-space within a shell and across shells have sharable information, which can be leveraged in a joint k-q reconstruction for accelerated dMRI. Antipodal symmetry property can be incorporated as q-space points at the opposite sides of the sphere share the same diffusion contrast. Blue and orange points refer to samples on different q-space shells. b) To enhance the joint k-q reconstruction, q-space neighborhood (across directions and shells) is designed to have highly diverse k-space undersampling patterns.

The GP covariance matrix **Σ** in Eq.1 is parametrized using a covariance function *C*. For multi-shell dMRI acquisition consisting of multiple b values (and directions), a valid covariance function between two q-space locations (*q, q*′) can be defined as:

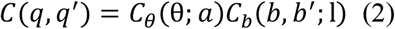

where *q* and *q*′ refer to q-space locations at two b shells (*b* and *b*′) with an angular distance of θ, *C*_0_ is a spherical covariance function describing relations between dMRI data within a shell and *C*_b_ is a squared-exponential function describing relations of dMRI data between shells [40]:

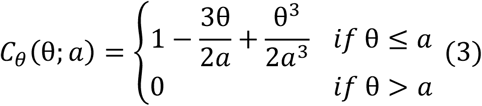

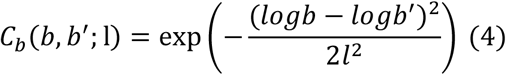

Where *a* is a hyperparameter controlling signal smoothness within shell and serves as an angular threshold, *l* is a hyperparameter controlling signal smoothness across shells.

Because dMRI measurements reflect both true signal variation and noise, two more hyperparameters are included in the covariance function to represent noise 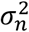 and signal variation *λ*. The covariance matrix can be calculated as:

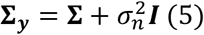

with **Σ**_i,j_ = λ(q_i_, q_j_), where **Σ**_***y***_ denotes the covariance matrix for noisy dMRI data **y**, and ***I*** is an identity matrix. In multi-shell condition, different 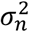 are estimated for different shells.

### GP estimation of dMRI signals

GP modelling allows us to construct a prior distribution of dMRI signals in q-space. Based on Bayes theorem, a posterior distribution can be calculated by updating the prior distribution with the likelihood from the data. The Maximum A Posteriori (MAP) probability provides an estimate of q-space signal based on this posterior distribution [41]:

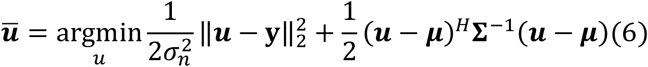

Where ū is the MAP point estimate, ***y*** is the noisy data.

### Multi-shell-DAGER reconstruction

Inspired by the use of GPs in dMRI pre-processing, DAGER incorporates GPs into the earlier step of image reconstruction and extends it to multi-coil acquisitions. DAGER uses the GP-estimated dMRI signal as a prior and solves the following reconstruction problem:

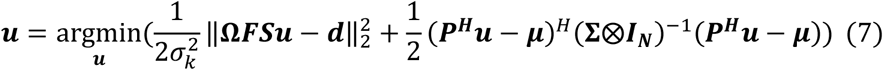

where ***u*** is the unknown image, **Ω** refers to undersampling operator, ***F*** refers to Fourier transform, ***S*** refers to coil sensitivity encoding, 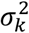 is the noise variance in the k-space data, which can be calculated from 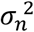 as in [35], ***d*** is the acquired k-space signal, *μ* is the mean value of GP prediction, **Σ** is the GP covariance matrix and *N* is the number of voxels, *H* is the conjugate operation and ⨂ is Kronecker product. ***p*** represents the motion induced phase error, which is different between diffusion directions. DAGER iteratively solves hyperparameter estimation and image reconstruction [35].

In this work, we extend DAGER to incorporate multi-shell information sharing in the reconstruction framework, resulting in ms-DAGER. The original single-shell covariance matrix **Σ** in Eq.7 is replaced with the multi-shell covariance matrix **Σ** defined in Eq.2.

### Correction for Rician noise fitting

The Gaussian Process assumes Gaussian noise in the data. However, in practice GP fitting is performed on magnitude images, where noise follows a Rician distribution. In scenarios where the noise level is high, the noise variance 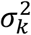 in Eq.7 estimated from GP model can deviate significantly from the true noise variance, impacting the accuracy of GP fitting.

To mitigate this issue, we propose to incorporate the noise estimation from the variance-stabilizing transformation (VST) method to improve the noise estimation accuracy. The variance-stabilizing transformation defines a nonlinear function *f*(*z*) to map the Rician random variable *z* to new values with variances that are independent of signal intensity (stabilized). The nonlinear mapping function *f*(*z*) and noise estimation are jointly optimized by minimizing a loss function consisting of a main term controlling the accuracy of the transformation (variance becomes independent of signal intensity) along with penalty terms to enhance the robustness of the estimation process [37].

## Method

### Simulation

We first evaluated the performance of ms-DAGER using realistic simulations based on HCP dMRI data consisting of 3 shells (1000, 2000 and 3000 s/mm^2^) and 270 directions[42]. The data was fitted with the ball-and-stick model which was used to generate simulation data with the same model[43]. A 100-direction dataset was generated with 50 b=1000s/mm^2^ and 50 b=2000s/mm^2^ directions uniformly sampled in q-space, which was used as a reference. Datasets with 36 and 64 directions were simulated by uniformly undersampling the 100-direction dataset. For ms-DAGER reconstruction, we optimized k-q undersampling using a graph model [44] that aims for each q-space neighbourhood (across directions and shells) to have highly diverse k-space undersampling patterns (Figure 1b). Undersampled multi-channel datasets were simulated with sensitivity maps from an 8-channel head coil and a total undersampling factor of 12 (in-plane R =3 and through-plane SMS=4) with added noise. This simulation intends to deliberately test ms-DAGER’s ability to exceed the theoretical limits by using a total acceleration that exceeds coil count.

To evaluate potential angular smoothing effects introduced by the Gaussian Process prior, we calculated the covariance between directions as a function of angular distance. We also calculated the point-by-point intensity ratio between direction-averaged images of different shells to evaluate the cross-shell smoothness.

### In vivo experiment

Seven subjects were scanned on a Siemens 7T scanner. Informed consent in accordance with local ethics was obtained before each scan. A 2D spin-echo dMRI sequence was modified to include SMS acquisition and blipped-CAIPI encoding with highly variable k-space undersampling patterns for each q-space neighbourhood. A 32-channel coil was used with coil sensitivities measured using a GRE sequence. The dMRI data were acquired with in-plane acceleration R=3 and multi-slice acceleration SMS=4 (total acceleration factor = 12). Three subjects were scanned with 1.25 mm isotropic resolution (TE=70ms) and four were scanned with 1.5 mm isotropic resolution (TE=68 ms). Other parameters: TR=3500ms, partial Fourier 6/8, 84 slices over 21 multi-slice sets.

The protocols were designed to evaluate the performance of ms-DAGER with a reduced number of diffusion directions (with fixed resolution) and different resolutions (with fixed diffusion directions). For the 1.5mm protocol, we acquired three dMRI datasets with 100, 66 and 36 diffusion directions staggered across two shells (b=1000s/mm^2^ and 2000s/mm^2^) using optimized k-q sampling patterns and equal number of directions for each shell (50 b=1000s/mm^2^ and 50 b=2000s/mm^2^, etc.). The q-space points for the 66-direction dataset were obtained by optimally undersampling those of 100-direction dataset, while the q-space points of the 36-direction dataset were separately designed for uniform angular coverage. The 1.25mm dataset was acquired with the same q-space sampling except for omitting the 36 direction set. A 3-shell in vivo dataset at 1.5mm resolution was also acquired for subject 1 with 40 diffusion directions for each b-shells across b = 1000/2000/3000 s/mm^2^ to evaluate the performance for more shells (same parameters as above except TE = 78ms). Seven b=0 volumes were also acquired, including one volume acquired with reversed phase-encoding direction for distortion correction with the FSL’s topup [45-46].

For each subject, we acquired a high-SNR reference consisting of two repetitions of non-slice-accelerated (“single-band”) data with matched TR (covering 21 slices to reduce scan time) and in-plane acceleration. For one subject, a whole-brain single-band reference image was also acquired with one repetition.

To compare our method with more commonly used accelerations, we acquired SMS =2 whole-brain data, along with our proposed SMS =4 whole-brain data for comparison, single band data was also acquired as reference. To compare ms-DAGER method with the previous single-shell DAGER (ss-DAGER), an additional set of data were acquired from one subject using separately optimized k-space sampling patterns for each shell (which is optimal for single-shell DAGER) for a fair comparison with ms-DAGER using the same q-space sampling pattern. The details of the protocols are listed in Table.S1.

### Reconstruction implementation

The motion induced phase error ***p*** in Eq.7 was estimated using a method similar to MUSE [47]. For each diffusion direction, an initial SENSE reconstruction is performed and a smooth phase map is then estimated from the k-space center.

The dMRI signal is identical at opposite points on a q-space sphere, a property known as “antipodal symmetry”. We leverage this property during ss-DAGER and ms-DAGER reconstruction by extending the local neighborhood around each direction to include the region around its antipodal reflection, as shown in Figure.1b.

Since brain white matter and gray matter tissues have different diffusivity property, the hyperparameters were separately updated for these two types of tissues using gray matter and white matter masks generated using FA and MD thresholds (FA>0.2,MD>0.0003 mm^2^/s for white matter regions; FA<0.2, MD<0.0009 mm^2^/s for gray matter regions). In other parts inside the brain, we used the same hyperparameters as for gray matter.

In this work, we used VST to estimate the noise variance directly from magnitude data with the aim of improving the initial noise variance estimation for the GP fitting. To account for spatially varying noise amplification due to parallel imaging reconstruction, we first used VST to estimate the noise variance at each spatial location. Then we divided the estimated noise variance by the g-factor [12] at the corresponding location. Finally, the averaged noise variance from all spatial locations was used as an initial likelihood estimation in Gaussian Process parameter fitting.

### Post-processing

For the in vivo data, Gibbs ringing artifacts were removed using degibbs3D from mrtrix3 (https://www.mrtrix.org/) followed by correction for susceptibility-induced distortions, eddy currents and motion using FSL’s topup and eddy tools. DKI and NODDI metrics were calculated using DIPY (https://dipy.org/) and FSL [48], respectively. We also fitted the data using the ball and stick model with Bedpostx [49]. For one subject, we compared the whole brain tractography results between the reconstructed images and the single band whole brain reference images using autoPtx toolbox [50].

## Results

Figure 2 shows simulation results with 100, 64 and 36 directions. Compared to SENSE, ms-DAGER and original single-shell DAGER (ss-DAGER) provide improved image quality with significantly reduced noise level for all protocols. However, as the number of directions decreases (e.g., 64 and 36 directions), ss-DAGER suffers from increased image errors because fewer directions are available from ss-DAGER for pulling information in a given shell. In comparison, ms-DAGER provides relatively high image quality even with 36 directions due to its ability to draw information from multiple shells. This behavior is quantified with normalized Root-Mean-Square-Error (nRMSE) values, which increase rapidly for ss-DAGER as the number of diffusion directions decreases, but remain similar for ms-DAGER, demonstrating improved robustness of ms-DAGER.

**Fig 2.**
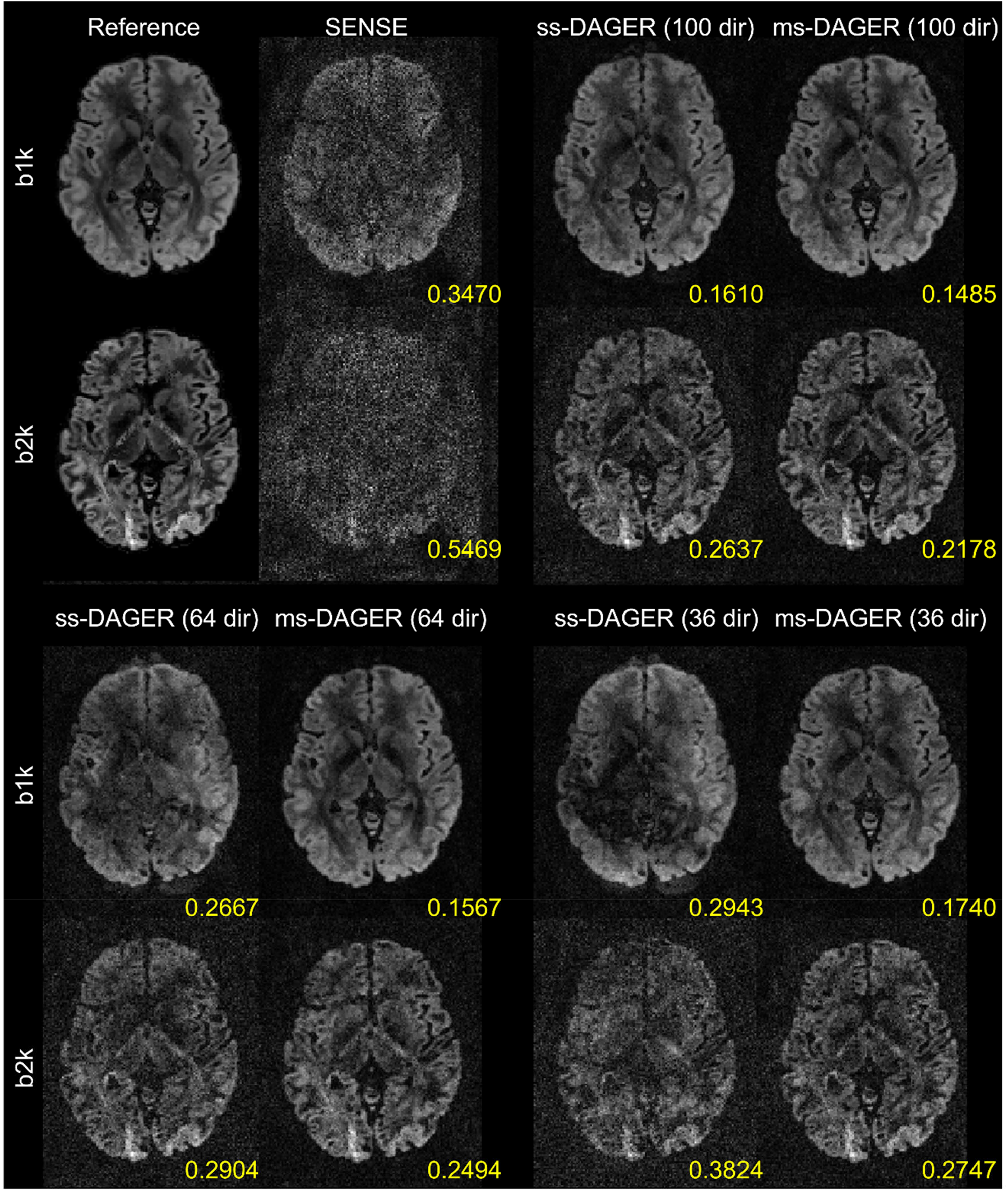
Reconstruction results of the 100/64/36-direction simulation data with undersampling factor R =12 (SMS=4, R=3). Reconstructed b=1000s/mm^2^ (‘b1k’) and b=2000s/mm^2^ (‘b2k’) images from SENSE, ss-DAGER, and ms-DAGER are shown for each method. Note that ms-DAGER produces the highest SNR and the least artifacts. Compared to ss-DAGER, ms-DAGER allows improved reconstruction by exploring shared information across shells, particularly with a small number of directions (i.e., 64 and 36 directions).

Because ms-DAGER leverages information from nearby directions and b-shells to improve the reconstruction, it risks introducing smoothness across both angle and shell. We investigated these two types of smoothness in ms-DAGER using simulation data. To investigate angular smoothness, angular covariance of the simulated ground truth dMRI signal and the ms-DAGER reconstruction were compared (Figure 3a). Because noise will decrease the covariance values and confound the smoothness evaluation, a noise-free simulation was used in this investigation. The covariance is plotted for all pairs of directions on the same b-shell as a function of their angular difference in q-space. ms-DAGER demonstrates highly consistent angular covariance profile for 100, 66 and 36 diffusion directions with respect to reference, indicating that ms-DAGER is introducing minimal angular smoothness. The discrepancy increases slightly at a low number of directions (e.g., 36-dir) because ms-DAGER has fewer points in a local neighborhood to exploit information redundancy. Cross-shell smoothness was evaluated by assessing the intensity ratio between different shells (Figure 3b). We calculated the point-by-point intensity ratio between direction-averaged b=1000 s/mm^2^ images and direction-averaged b=2000 s/mm^2^ images for the simulated ground truth and ms-DAGER reconstruction within a white matter mask. Histograms are shown for two different noise levels (SNR 20 and 40 based on fully sampled data) and different numbers of diffusion directions. Cross-shell smoothness would lead to reduced ratio compared to the ground truth. The intensity ratio between two shells was slightly decreased with ms-DAGER (3.8% at most, for 36 directions), indicating minimal cross-shell smoothing.

**Fig 3.**
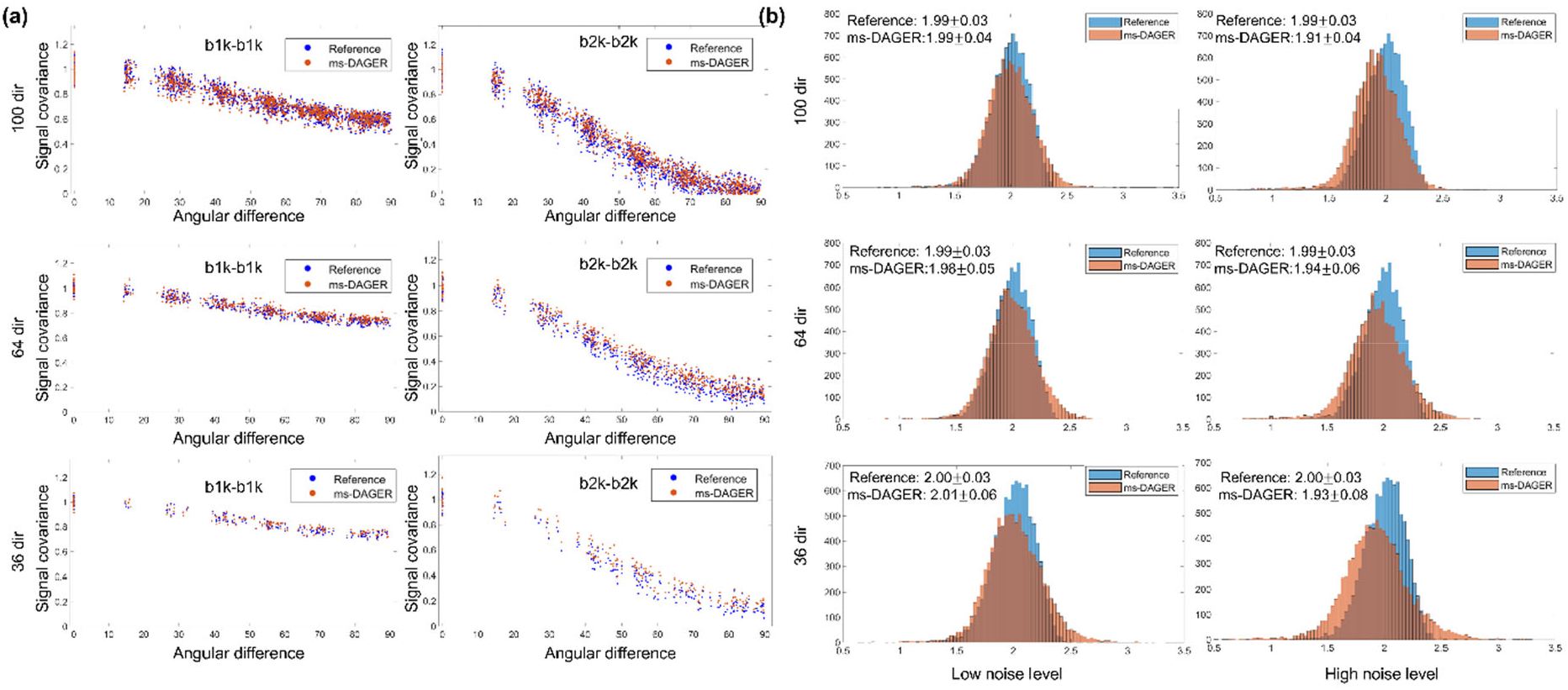
(a) Signal covariance of dMRI images. Whole-brain white matter of the simulation data was used for the covariance calculation. Each point represents 1 diffusion direction. For each data set, the covariance is normalized by the median of signal variances (angular distance = 0°). ms-DAGER reconstructed images demonstrate consistent covariance with the reference at both b=1000s/mm^2^ (‘b1k’) and b=2000s/mm^2^ (‘b2k’) shells. (b) Histogram of point-by-point intensity ratio between direction-averaged b1k images and direction-averaged b2k images, calculated with whole-brain white matter signal of the simulation data. High noise level and low noise level refers to SNR=20 and SNR=40 data (SNR calculated based on fully sampled data).

In Figure 4, we compared the noise estimation accuracy and effects on reconstruction with and without using VST by adding different levels of noise to simulation data (noise standard deviation 1-16 refers to SNR 30 to 1.875). As shown in Fig.4a, with the increasing of noise levels added to the simulation, the results without using VST (w/o VST) show strong negative bias, which increases with the ground truth noise standard deviation, revealing a large underestimation effect at high noise levels. In comparison, VST based method (w/ VST) provides a more robust estimation of noise standard deviation. As shown in Fig.4(b), the nRMSE values between reconstruction results and the ground truth at different noise levels are decreased with the introduction of VST.

**Fig 4.**
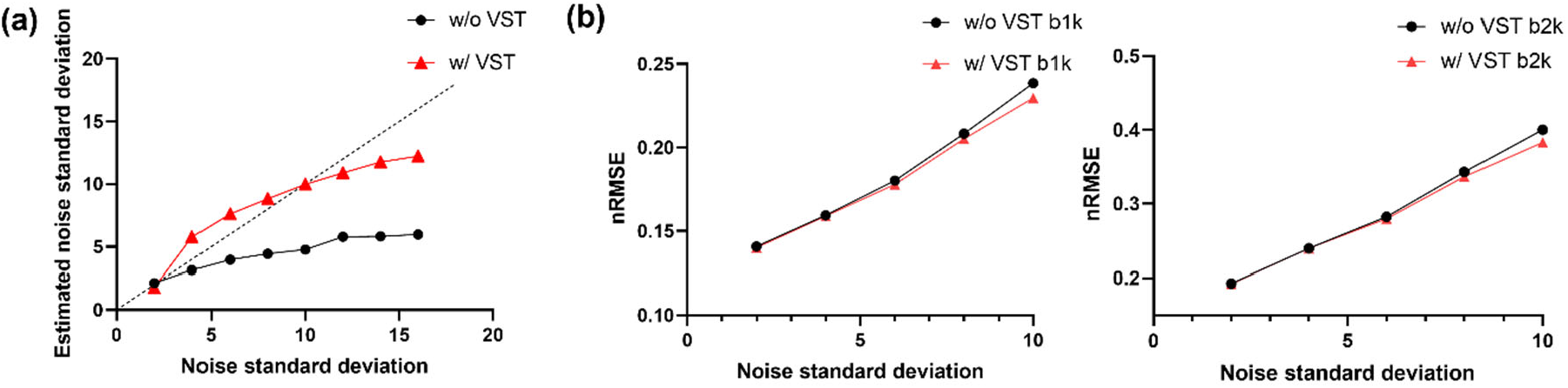
(a) Comparison of noise standard deviation estimation for the simulation data. Different noise levels were added to the ground truth. The noise estimation with VST (w/ VST) and without VST (w/o VST) are shown. VST allows more robust estimation of noise standard deviation, particularly at high the noise levels. (b) nRMSE values between ground truth and reconstruction results with and without using VST for noise standard deviations of 2, 4, 6, 8, 10. With the increase of noise levels, the method using VST shows better performance compared to the method without using VST.

Figure 5 shows the reconstruction results for the 1.5 mm in vivo data from subject 1 with 36-directions using SENSE, ss-DAGER and ms-DAGER. The k-space undersampling patterns are separately optimized for ss-DAGER and ms-DAGER (the q-space sampling patterns are identical). Compared to SENSE, both ss-DAGER and ms-DAGER methods significantly improve the image quality. However, the ss-DAGER reconstruction is noisier than ms-DAGER, particularly for the b = 2000 s/mm^2^ shell, whereas ms-DAGER provides similar data quality to the high-SNR single band reference (2 averages). These results demonstrate the advantage of ms-DAGER over original DAGER when the number of diffusion directions is small.

**Fig 5.**
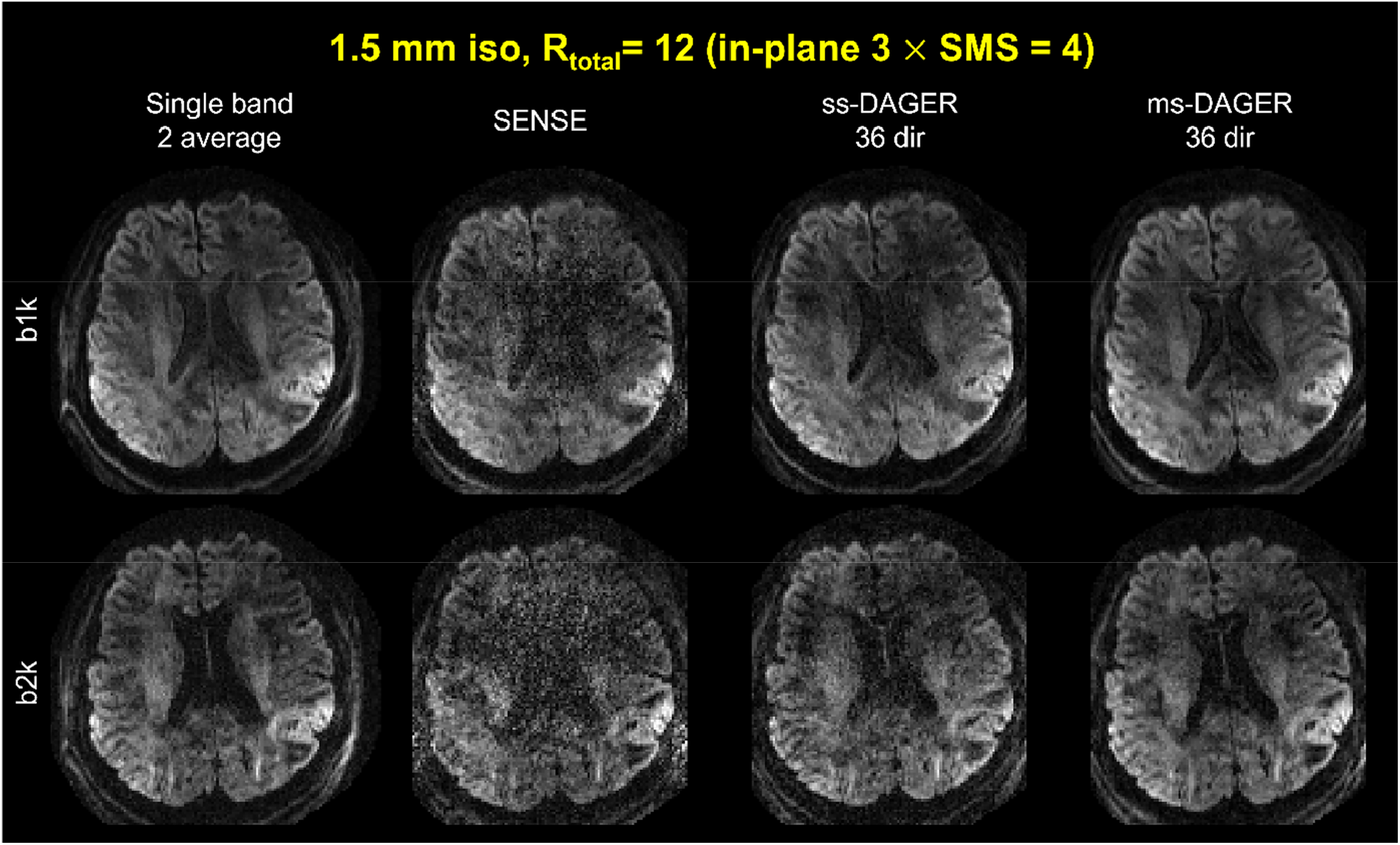
Reconstruction results of the 36-direction 1.5 mm isotropic resolution in vivo data. Single band reference images with 2 averages, SENSE, single shell DAGER (ss-DAGER) and ms-DAGER results at b=1000s/mm^2^ (‘b1k’) and b=2000s/mm^2^ (‘b2k’) are shown. Compared to ss-DAGER, ms-DAGER further improves the image quality when the number of diffusion directions is small.

Figure 6 shows an axial slice of the reconstruction results for the 1.5 mm in vivo data from subject 2 with 100, 66 and 36 directions. Compared to the SENSE results, ms-DAGER provides significantly improved SNR and reduced artifacts, with consistent structure details to the high SNR single-band reference. Even with a small number of diffusion directions as 36 directions, the results still show comparable quality to the high SNR reference.

**Fig 6.**
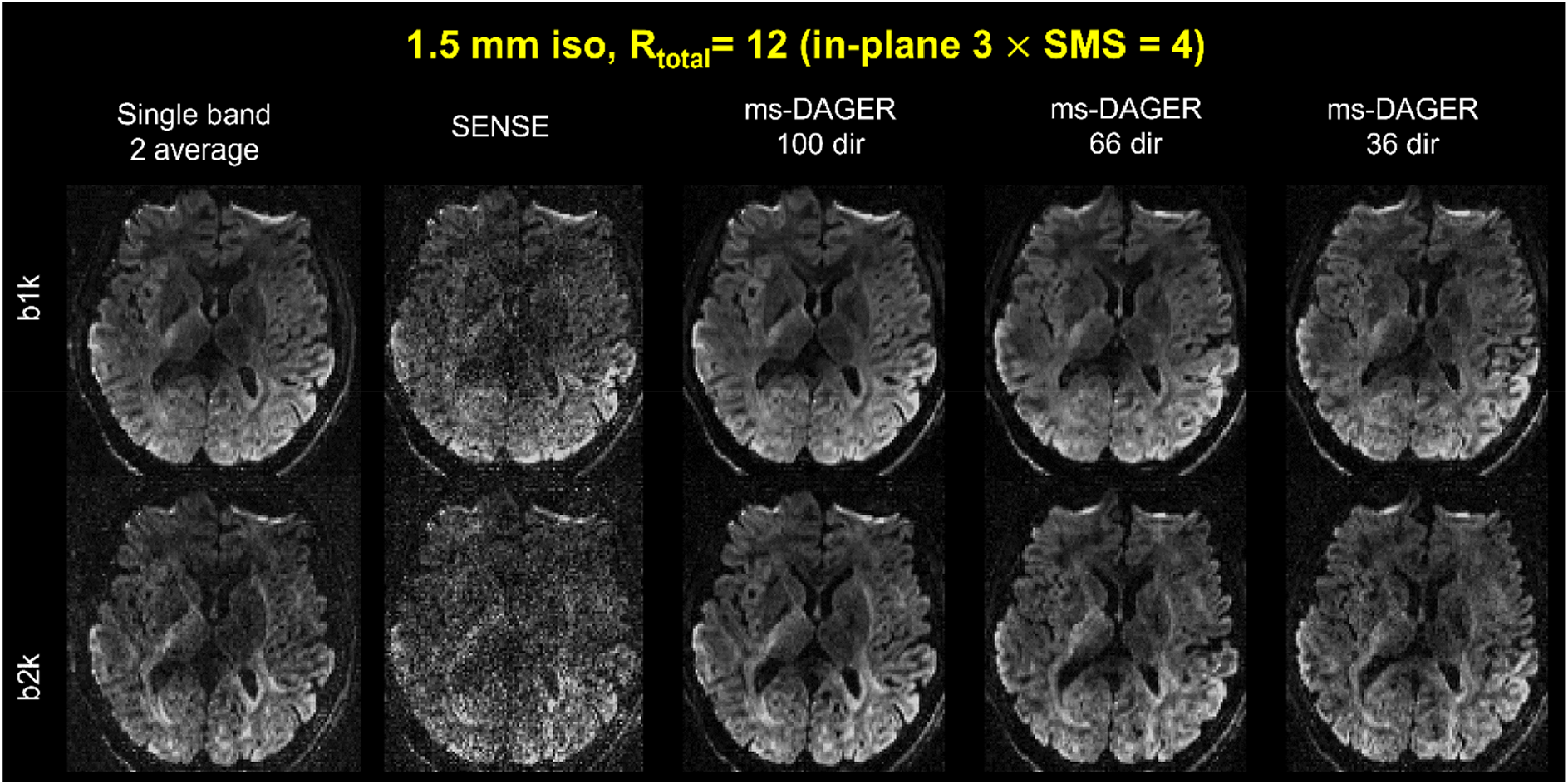
1.5 mm isotropic resolution in vivo data with different number of directions. Single band reference images with 2 averages, SENSE and ms-DAGER with different number of directions are shown at b=1000s/mm^2^ (‘b1k’) and b=2000s/mm^2^ (‘b2k’). ms-DAGER improves image quality compared to SENSE and provides consistent detailed structural information with the reference images.

ms-DAGER reconstruction results for two other subjects (1 and 4) with 1.5mm isotropic resolution are shown in Supporting Figure S1. ms-DAGER consistently improves the image quality compared to SENSE and shows comparable data quality to the high SNR reference images for different numbers of diffusion directions, which demonstrates high reproducibility of ms-DAGER.

In Supporting Figure S2, we compare our method with a SMS acceleration factor of 2, as is currently common practice, along with a single band whole-brain reference from subject 3. Sagittal, and coronal views are shown. Here, the single band reference was acquired with only one average due to scan time constraints. All data were acquired with the same in-plane acceleration factor of 3. As shown in Supporting Figure S2, the 12-fold acceleration data reconstructed using ms-DAGER show comparable image quality to the 3-fold accelerated single band reference and improved SNR compared to the 6-fold accelerated SMS=2 data, particularly for the higher b value shell. Note the image contrast is slightly different between protocols due to TR difference.

Figure 7 shows the reconstruction results for the 1.25mm in vivo data for subject 5. The image quality is extremely poor in SENSE reconstruction, which is dominated by noise. ms-DAGER significantly improves the image quality for 100-dir data and 66-dir data, showing consistent structure details compared to reference.

**Fig 7.**
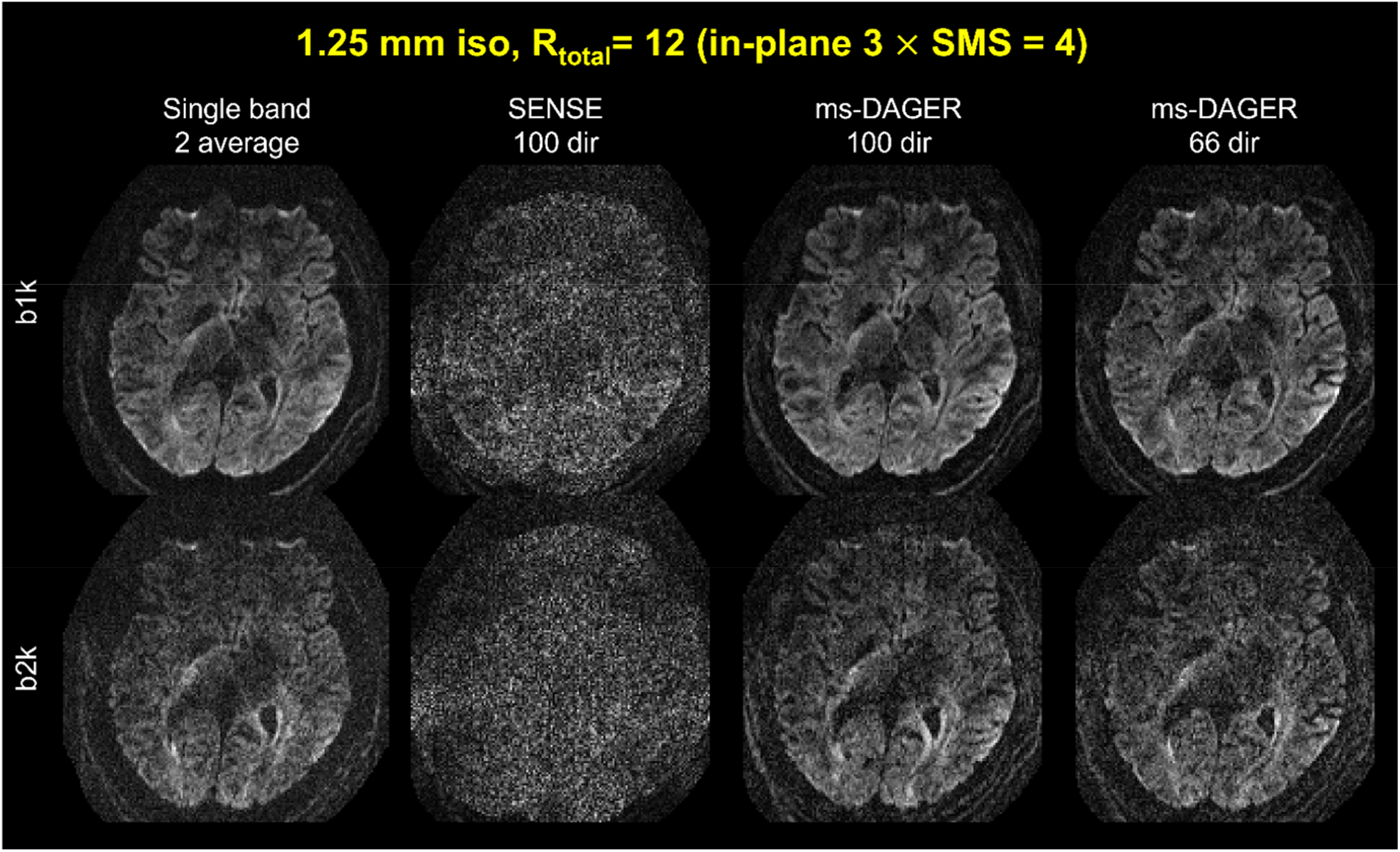
Reconstruction results for the 1.25 mm isotropic resolution in vivo data. Single band reference images with 2 average, SENSE and ms-DAGER results for different number of diffusion directions are shown. b=1000s/mm^2^ (‘b1k’) and b=2000s/mm^2^ (‘b2k’) images are both shown. Despite low SNR at 1.25mm resolution, ms-DAGER provides robust reconstruction from highly-undersampled multi-shell datasets.

DKI fitting results, including mean kurtosis (MK), axial kurtosis (AK) and radial kurtosis (RK) are demonstrated in Figure 8, SENSE reconstructions have inflated MK and AK values compared to the reference while ms-DAGER produces results consistent with the reference for matched number of directions. As the number of directions reduces, DKI maps with ms-DAGER have more outliers (i.e., black dots in the maps, indicating fitting failures), but the overall spatial patterns are consistent with the high-SNR reference.

**Fig 8.**
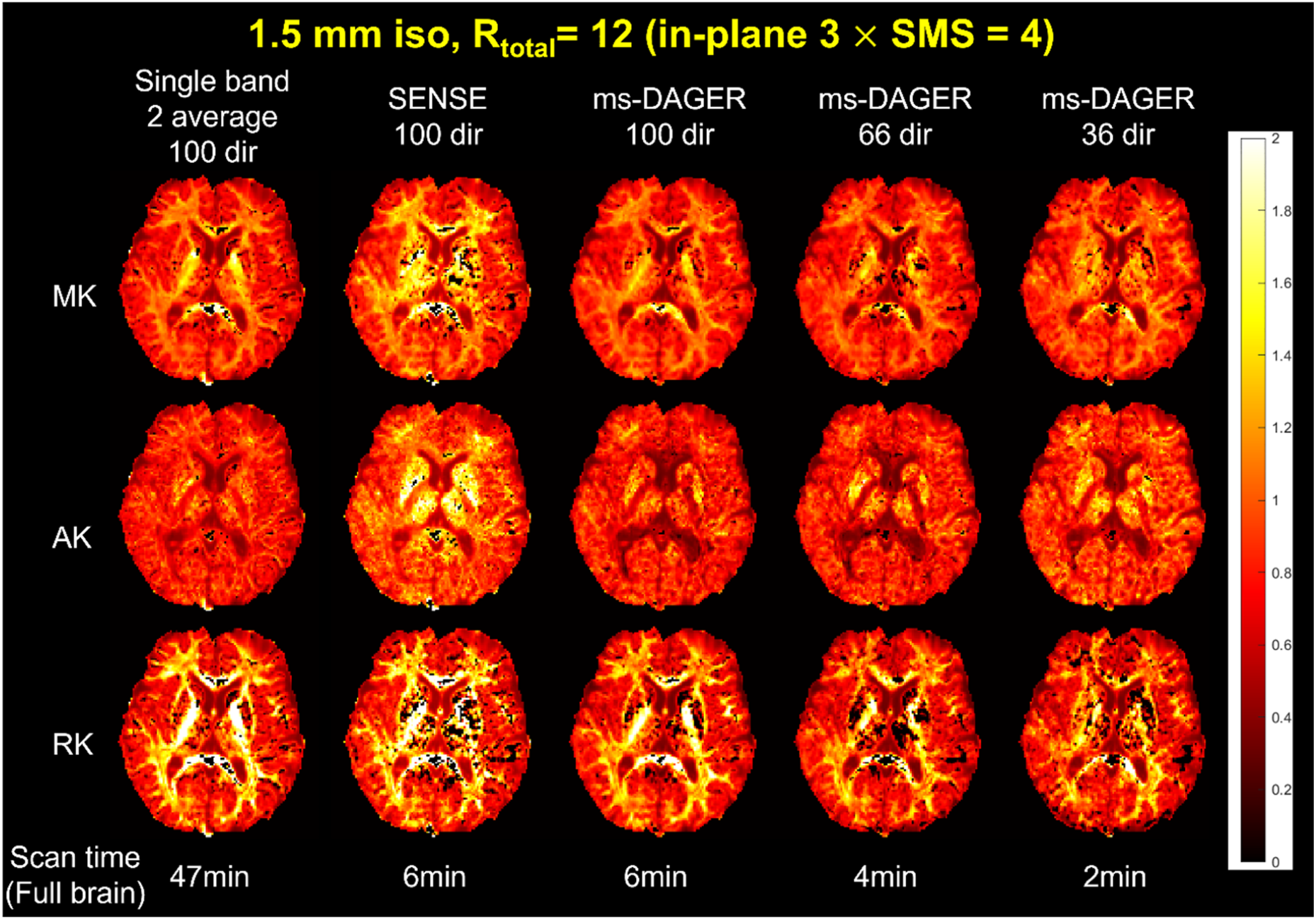
DKI fitting results for the 1.5 mm isotropic resolution in vivo data. Mean kurtosis (MK), axial kurtosis(AK) and radial kurtosis(RK) maps are shown calculated from single band reference images with 2 average, SENSE and ms-DAGER for different numbers of diffusion directions. ms-DAGER accelerated reconstruction generates similar DKI metric maps as the 2 average single-band reference with a much shorter scan time.

Figure 9 compares NODDI modelling fitting results from subject 2, specifically maps for isotropic fraction (fiso), intra-cellular fraction (fintra), and orientation dispersion index (ODI). ms-DAGER produces parameter maps that are consistent with the high-SNR reference while providing full brain coverage with a much shorter scan time. The degraded data quality in SENSE reconstruction (e.g., in Figures, 5 and 6) leads to instability in the modelling fitting, including overestimated fintra and large errors in thalamus (white arrow) for ODI due to the high noise level. In comparison, ms-DAGER provides parameter maps that are consistent with the reference with matched number of directions. When the number of directions reduces, ms-DAGER provides robust NODDI fitting despite significantly reduced scan time (e.g., 2min for 36-dir ms-DAGER versus 47min for 100-dir reference).

**Fig 9.**
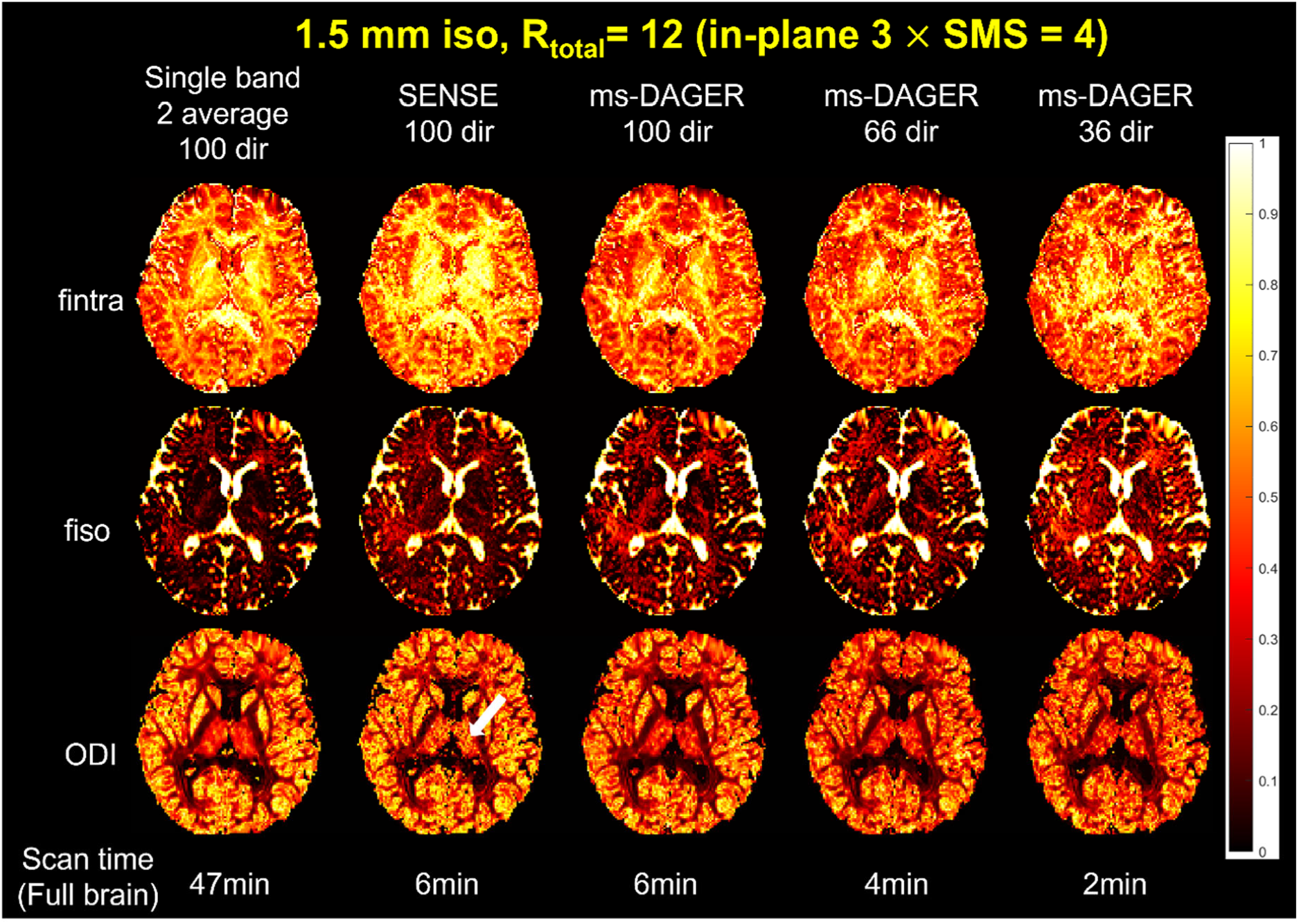
NODDI fitting results for the 1.5 mm isotropic resolution in vivo data. CSF volume fraction(fiso), intra-cellular volume fraction(fintra) and orientation dispersion index (ODI) from NODDI fitting are shown, which were calculated from single band reference images with 2 average, SENSE and ms-DAGER results for different numbers of diffusion directions. ms-DAGER accelerated reconstruction produces comparable NODDI metric maps as the single-band reference with a much shorter scan time.

Supporting Figure S3 shows the ball and stick model fitting results using FSL’s Bedpostx tool for the 1.5mm in vivo data from Subject 2, including both primary and secondary fiber populations. With 100 diffusion directions, both SENSE and ms-DAGER capture similar primary fiber populations. However, due to the high noise level in the SENSE reconstruction, the number of voxels with a robust secondary fiber population is significantly decreased as Bedpostx filters out secondary fiber population if they are unsupported by the data due to low SNR. In comparison, ms-DAGER results identify a secondary population in many more voxels, which have consistent patterns to the reference. As shown in the images, the orientations of secondary fibers are also consistent with the high SNR reference results.

Figure 10 shows tractography results for subject 3. autoPtx is used to track the fibers for 66-dir whole-brain single band reference data (16min) along with SMS 4 66-dir whole-brain data (4min) reconstructed using SENSE and ms-DAGER. Most fiber bundles are properly delineated from the single band reference data. Compared to the reference, many tracts are either incomplete or missing when using SMS 4 SENSE reconstructions for example acoustic radiation tracts (Figure10, middle column, in green), forceps minor (Figure10, left column, in yellow), inferior frontal-occipital fasciculus (Figure10, right column, in green) are absent. ms-DAGER reconstructions capture most tracts, consistent with single band reference with a much shorter scan time.

**Fig 10.**
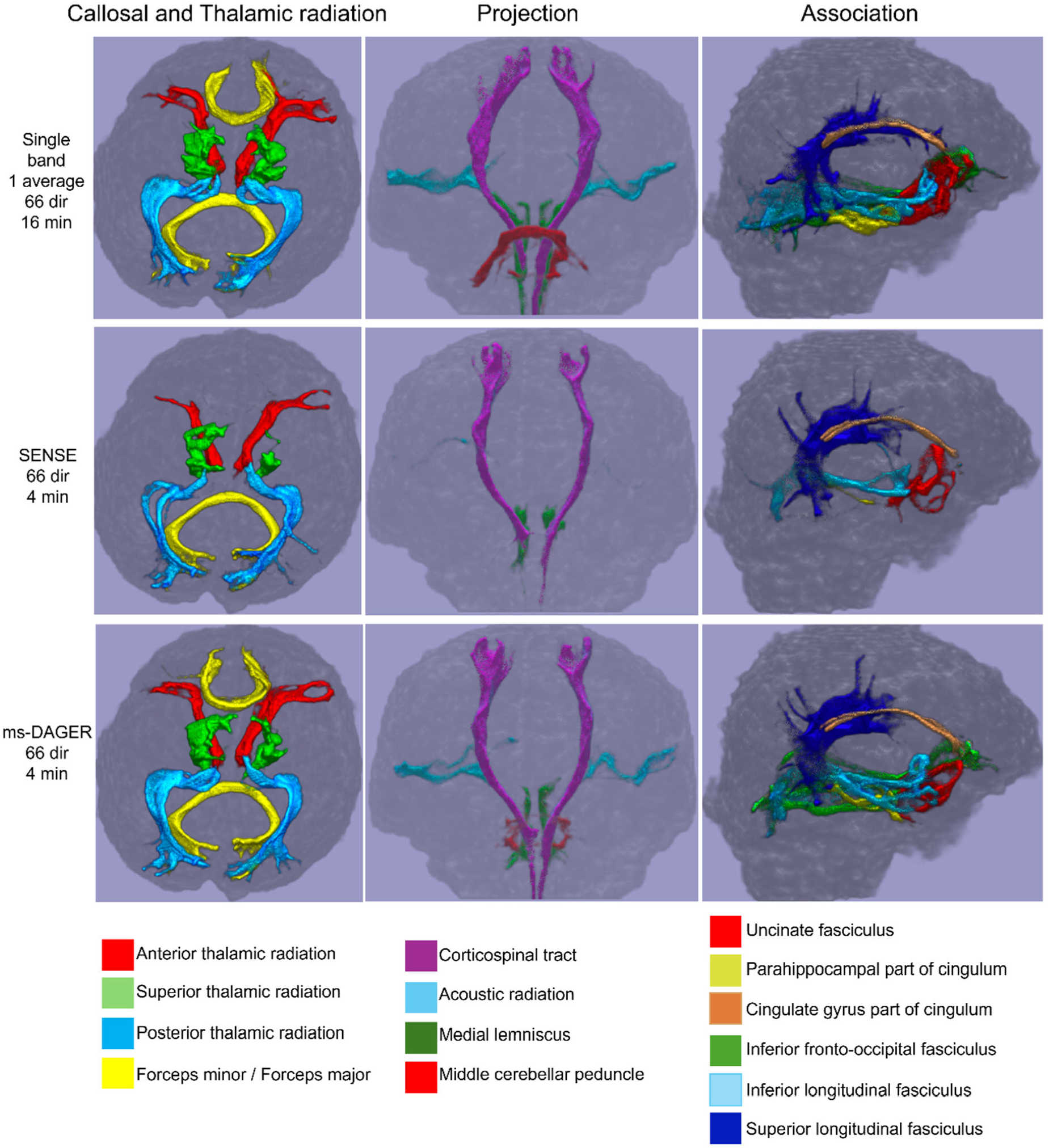
Fiber tracking results for the 1.5 mm isotropic resolution in vivo data using autoPtx. Single band reference, SENSE and ms-DAGER for SMS= 4 data are shown respectively. 14 major white matter pathways (constituting 27 separate tracts in right and left hemispheres) are rendered in superior (thalamic radiation and callosal fibers), anterior (projection) and lateral (association fibers) views. Compared to Single band, several fibers are totally missing in the SENSE results due to the high noise level while ms-DAGER manages to recover most of the fiber tracts.

## Discussion

In this work, we propose a robust acceleration method for multi-shell dMRI to improve acquisition efficiency while preserving image fidelity. This method builds on our previous single-shell DAGER approach, using Gaussian processes to model smoothness of the dMRI signal in q-space neighborhoods and leverage the associated covariance structure in image reconstruction. Simulation and in vivo results demonstrate that the proposed method can effectively preserve the SNR and suppress artifacts even at a high acceleration factor of 12 (SMS=4 and in-plane R=3). Unlike single-shell DAGER, ms-DAGER jointly reconstructs multiple shells to exploit information redundancy both within and across shells, allowing more efficient information sharing. To achieve robust model parameter estimation at low SNR, which is common to high-b values and high resolution dMRI, we further integrate a variance-stabilizing transformation to improve the accuracy of noise variance estimation when optimizing Gaussian Processes hyperparameters.

One major concern for joint k-q reconstruction is that when sharing information across different q-space points, there might be extra smoothing introduced beyond what is intrinsic to the data. This could introduce bias to model parameters reflecting fiber dispersion if angular smoothness is introduced or metrics reflecting non-Gaussianity if cross-shell smoothness is introduced. In this study, we investigated within-shell angular and cross-shell smoothness using simulated datasets. As shown in Figure 3, the ms-DAGER reconstruction only slightly increases the covariance between different directions and affects intensity ratio between shells compared to the reference, indicating that Gaussian Process is able to learn and exploit the smoothness intrinsic to the data without introducing much smoothing.

Noise estimation plays a key role as a hyperparameter in GP fitting. In the context of ms-DAGER, the estimated noise level is directly used as a regularization factor on the data consistency term (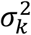 in Eq.7). Hence, estimating noise level accurately is crucial to reconstruction quality. In this work, we used VST to mitigate the noise variance estimation bias due to the Rician distribution in magnitude images when the noise level is high. We assessed the accuracy of noise estimation using simulated data with ground truth noise std from 1 to 10 in Fig.4. The results show that VST can improve noise estimation and hence improve image reconstruction accuracy.

One impact that ms-DAGER could have is to facilitate use of multi-shell dMRI models within clinically feasible scan times. Due to the noise amplification when reconstructing high acceleration factors, results from traditional parallel imaging methods like SENSE provide unreliable fits to models like NODDI or DKI, as shown in Figure 8 and Figure 9. ms-DAGER can significantly improve the image quality and maintain diffusion contrast information under these conditions of high acceleration, enabling more accurate results for both multi-shell model fitting and fiber tracking.

Supporting Figure S4 and S5 shows the DKI and NODDI fitting results of 1.25mm in vivo data of subject 5. In Figure S4, DKI fitting results from SENSE are corrupted by noise, showing large errors. In comparison, ms-DAGER produces more consistent results compared to the high SNR reference, though there are some outliers (black dots) due to the high noise level. As shown in Figure S5, SENSE results cannot support reliable NODDI fitting, leading to large errors in the parameter maps. In comparison, ms-DAGER allows better NODDI fitting even with a smaller number of diffusion directions of 66. The ODI parameters with ms-DAGER are slightly under-estimated compared to the high SNR reference, and the biases may be caused by the relatively low SNR at higher resolution. To investigate this effect, we also tested on other subjects. Supporting Figure S6 and S7 show the reconstructed images and NODDI fitting results for subjects 6 and 7 with 1.25 mm isotropic resolution, the model fitting results consistently show significant improvements compared to SENSE results and high reproducibility across subjects.

In this work, we primarily focused on dMRI protocols with two b-shells, which are commonly used in dMRI studies due to their compatibility with a large range of dMRI modelling fitting approaches and feasible acquisition times. We tested the performance of ms-DAGER with a set of 3-shell data (b=1000, 2000, 3000s/mm^2^). The preliminary results shown in Supporting Fig.S8 demonstrated that the image quality of higher b value shell can be improved by sharing information across shells even at b=3000s/mm^2^.

Our current study has some limitations. Firstly, the proposed method has not considered eddy currents. Eddy current effects can lead to geometric distortions and phase mismatch between diffusion directions and b values. The phase mismatch for different directions is captured in the phase error term ***p*** in Eq. 7, thus corrected during the reconstruction. For the distortions effect, we used a high in-plane acceleration factor R=3 in all our in vivo scans, resulting in modest eddy current induced distortions. We investigated this effect with a phantom scan using in-plane acceleration factor R=3 as used for in vivo scans with the identical EPI readout and calculated the between-volume voxel shift. The results showed a max shift of 0.292 pixel with a standard deviation of 0.234 pixel, which did not introduce a major bias in the joint k-q reconstructions performed in this work. Second, we have not accounted for potential subject motion, which will have similar (but less predictable) impact on the k-q framework. In the present work, we used experienced subjects who were instructed to remain as still as possible. In examining our SENSE results, we did not observe substantial motion in any of our subjects, which may in part reflect the relatively short scan times. The high quality of ms-DAGER results with crisp image edges also suggests negligible impact of eddy current distortions and motion. Nevertheless, eddy current induced distortions will be stronger with lower in-plane acceleration and motion will present in less compliant subjects. We will investigate the effects of the resulting mismatches and extend DAGER to include a correction framework in future work. Finally, the computation time of the current MATLAB implementation is long, which can be improved with algorithmic optimization, parallel computing, and more efficient implementation.

## Conclusion

We have proposed a novel reconstruction method based on Gaussian Processes to accelerate multi-shell dMRI data acquisition. Based on simulations and in vivo data, our proposed method enables multi-shell dMRI data with high spatial and angular resolution in shorter scan times compared to conventional methods. These data are of sufficient quality to support advanced modelling of microstructure and tractography.

## Acknowledgement

W.W. is supported by the Royal Academy of Engineering (RF\201819\18\92). K.L.M. is supported by the Wellcome Trust (WT202788/Z/16/A). This study is supported by the NIHR Oxford Health Biomedical Research Centre (NIHR203316). The views expressed are those of the author(s) and not necessarily those of the NIHR or the Department of Health and Social Care. The Wellcome Centre for Integrative Neuroimaging is supported by core funding from the Wellcome Trust (203139/Z/16/Z and 203139/A/16/Z). For the purpose of Open Access, the authors have applied a CC BY public copyright license to any Author Accepted Manuscript (AAM) version arising from this submission. The authors wish to thank Dr. Jesper Andersson for his help when using FSL’s eddy.

## Supporting information

**Table S1.**
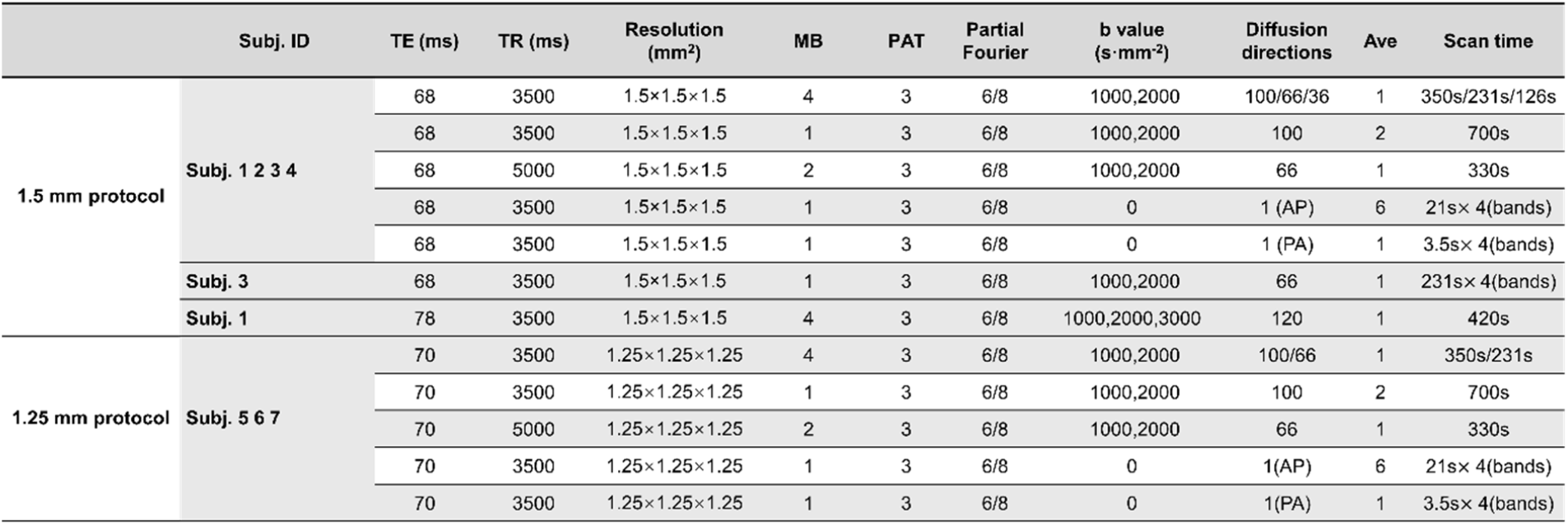
In vivo acquisition protocols.

**Fig S1.**
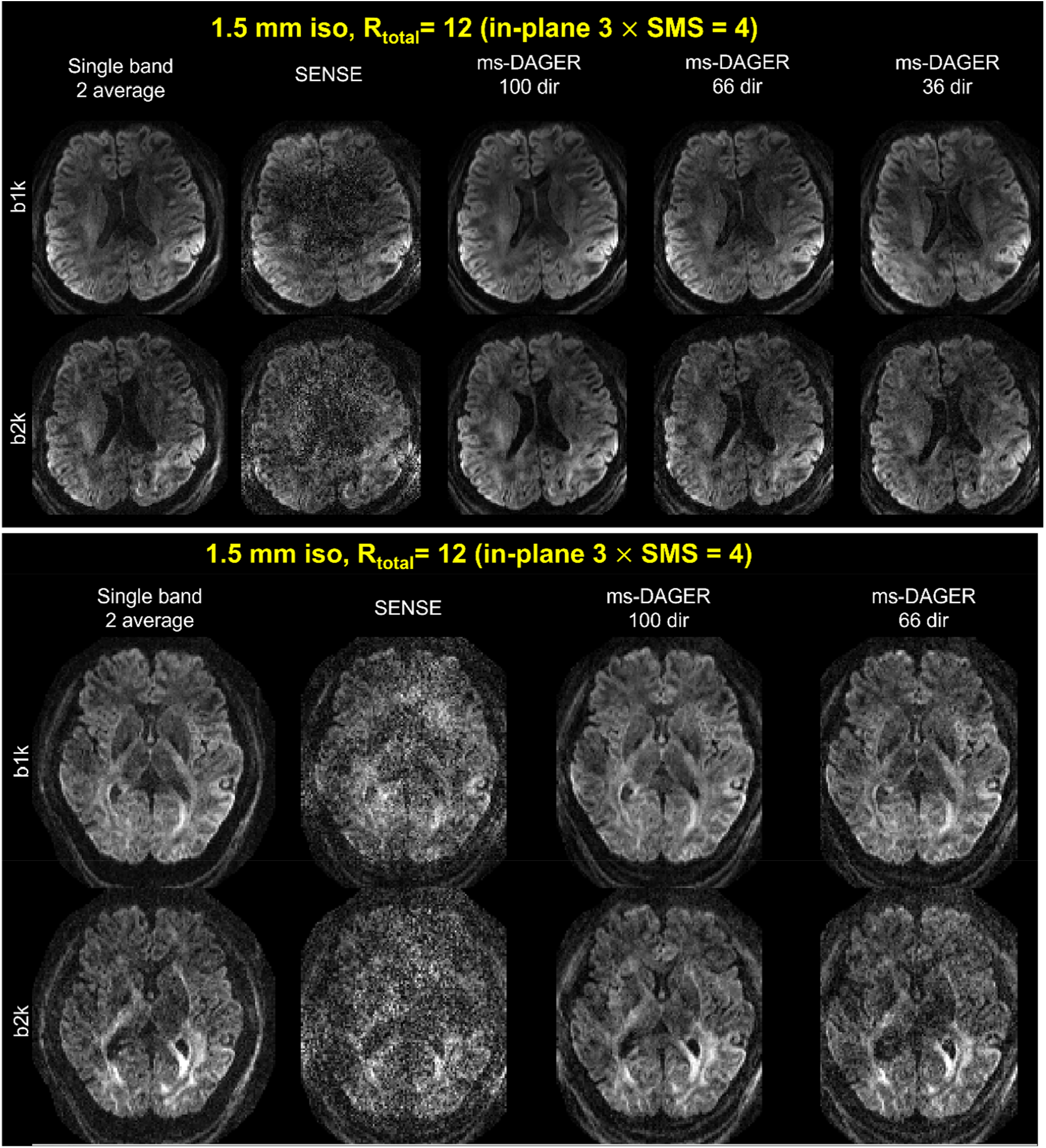
Reconstruction results of the 1.5 mm isotropic resolution in vivo data from two other subjects. Single band reference images with 2 average, SENSE and ms-DAGER results are shown b=1000s/mm^2^ (‘b1k’) and b=2000s/mm^2^ (‘b2k’) images are both shown. ms-DAGER consistently improve image quality compared to SENSE for both subjects.

**Fig S2.**
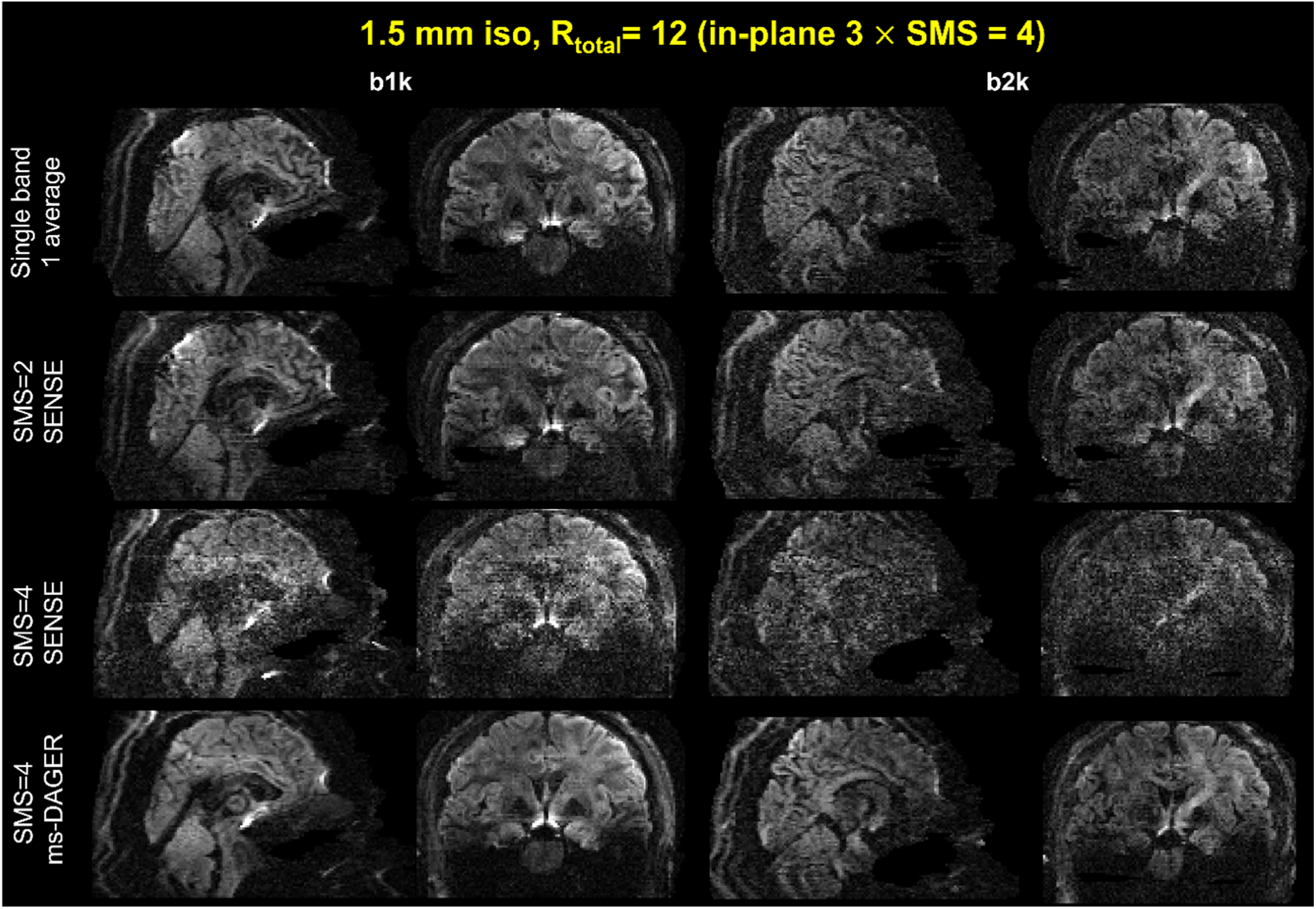
Coronal and sagittal slices of 1.5 mm isotropic resolution in vivo data. Single band reference images with 1 average, SMS=2 SENSE, SMS=4 SENSE and SMS=4 ms-DAGER results are compared. b=1000s/mm^2^ (‘b1k’) and b=2000s/mm^2^ (‘b2k’) images are both shown. ms-DAGER provides improved SNR compared to SMS=4 and SMS=2 SENSE, with comparable data quality to the reference images. Note image contrasts are slightly different between methods due to different TR used.

**Fig S3.**
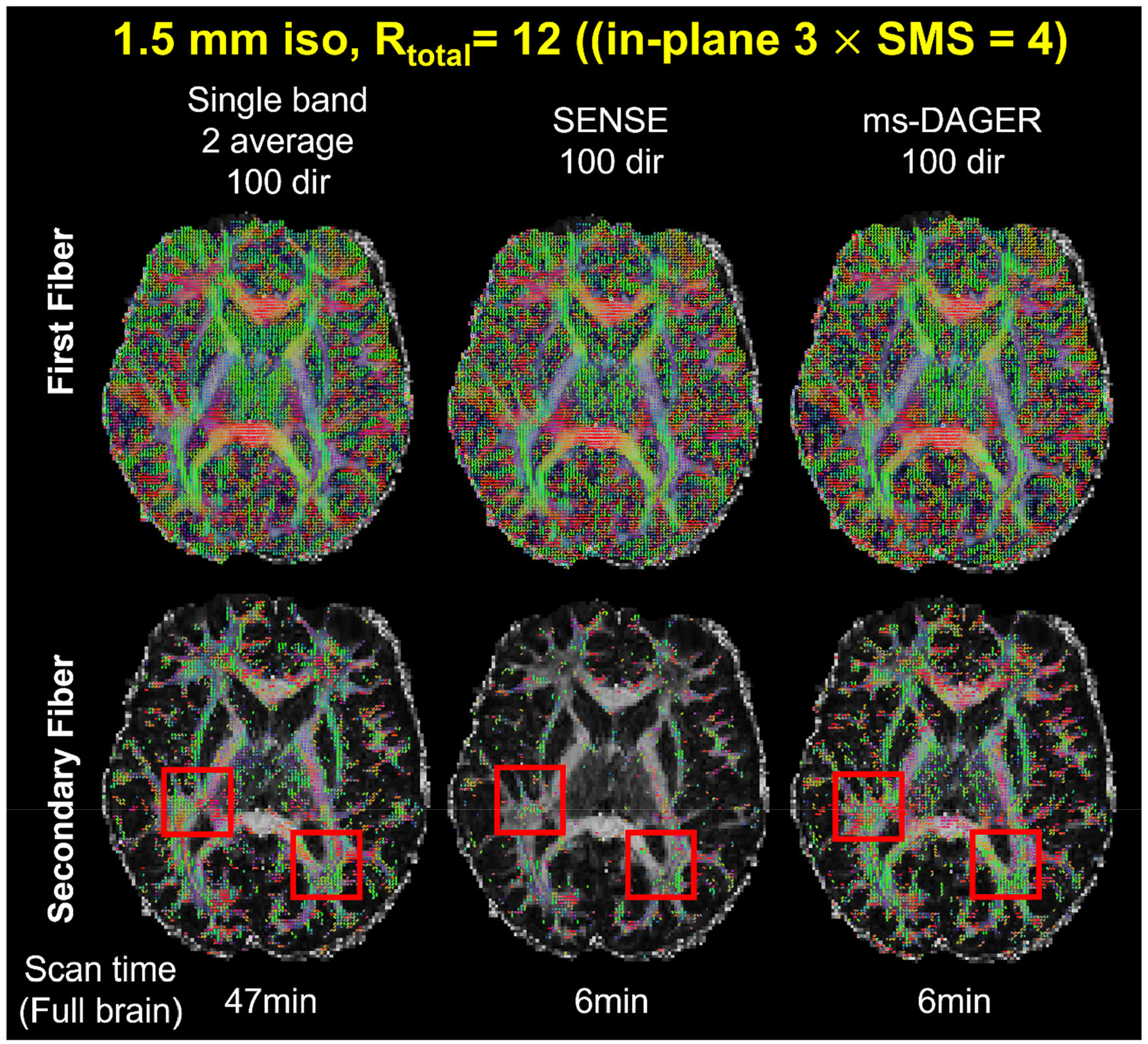
Ball and stick fitting results for the 1.5 mm isotropic resolution in vivo data. The first fiber and second fiber populations calculated from single band reference images with 2 average, SENSE and ms-DAGER results are shown. ms-DAGER recovers a large number of second fiber population, consistent with the single-band reference, while SENSE results fail to capture many second fibers.

**Fig S4.**
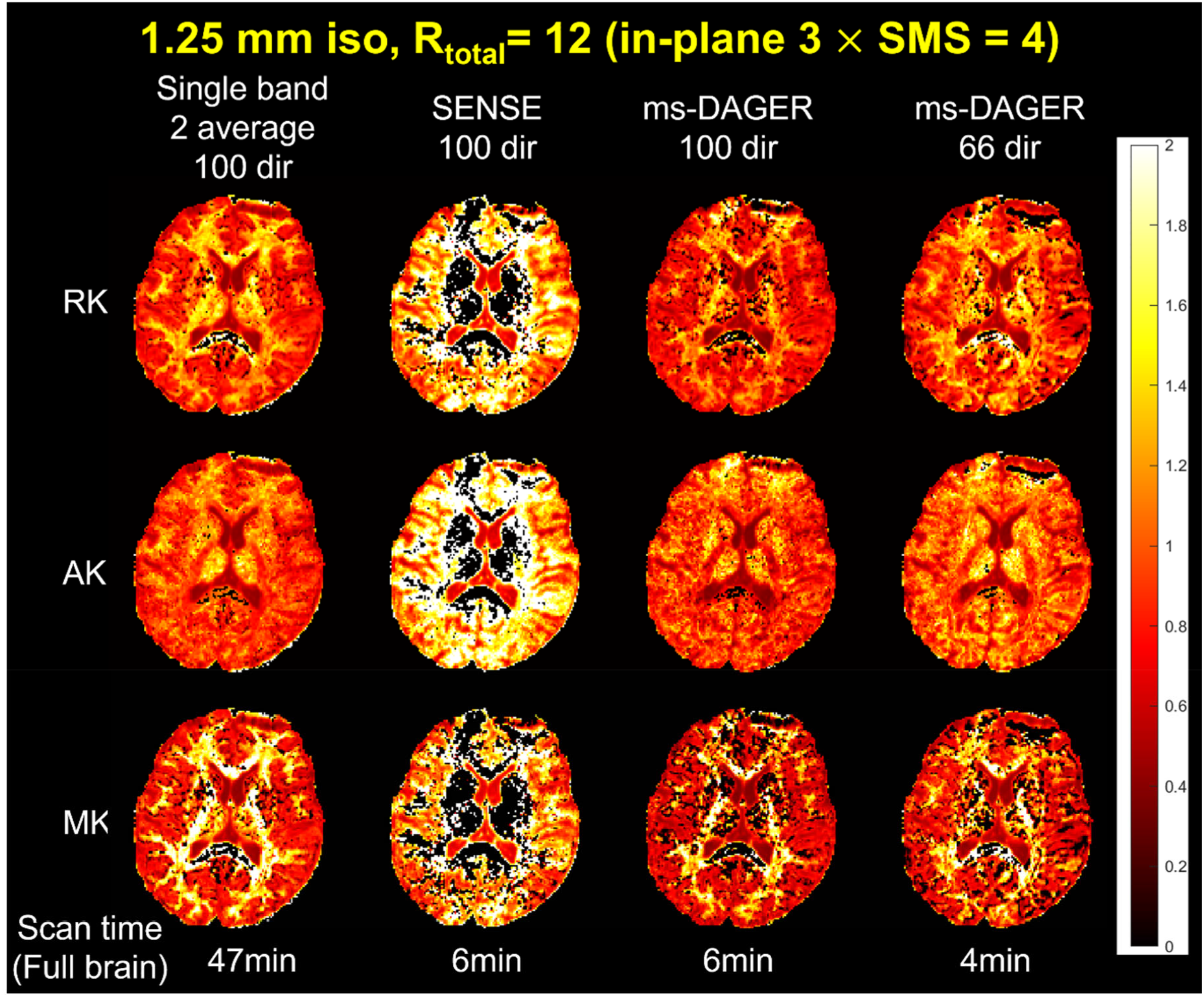
DKI fitting results for the 1.25 mm isotropic resolution in vivo data. Mean kurtosis (MK), Axial kurtosis(AK) and Radial kurtosis(RK) maps calculated from Single band reference images with 2 average, SENSE and ms-DAGER results are shown. Compared to SENSE, ms-DAGER produces more consistent results with reference using much shorter scan time.

**Fig S5.**
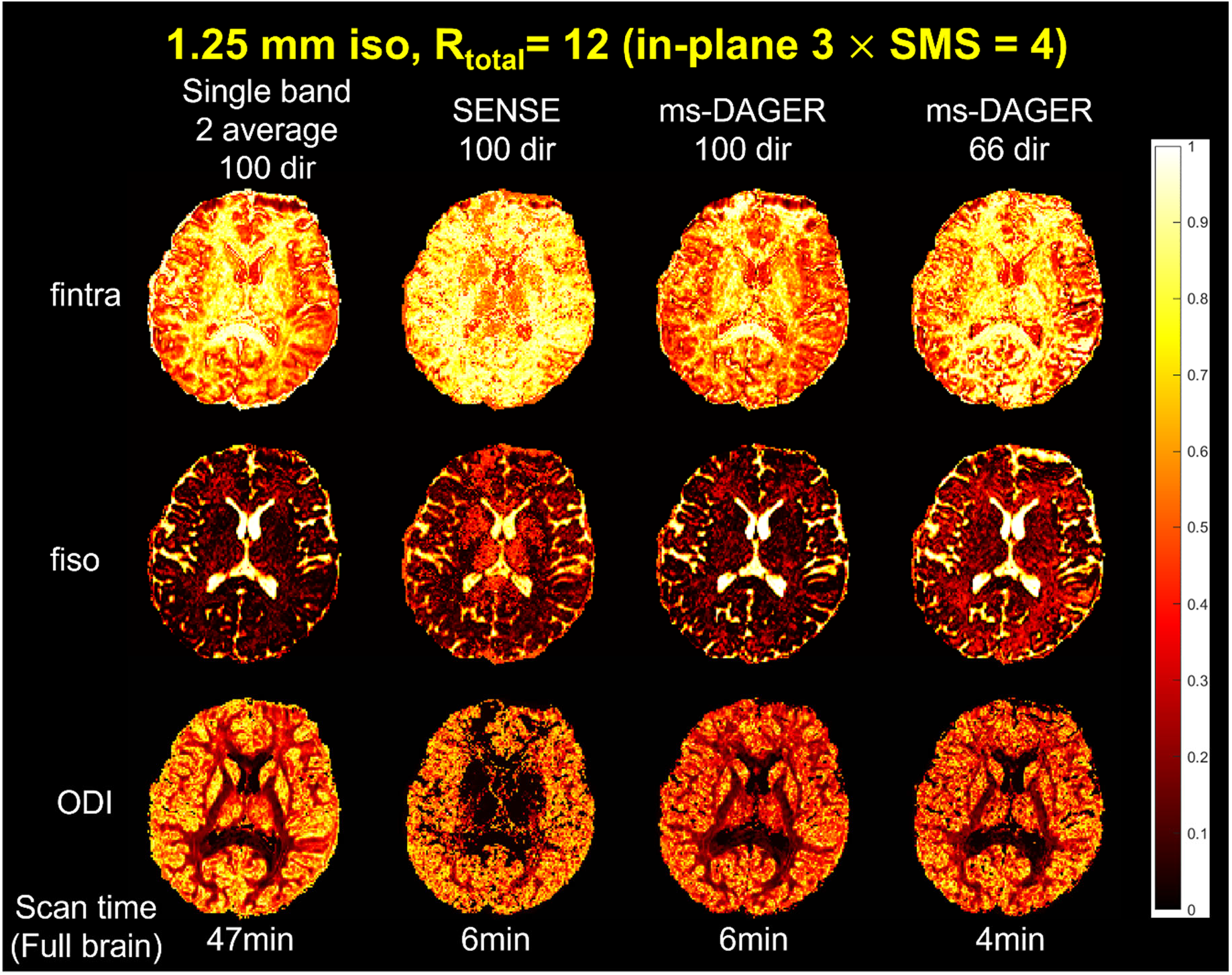
NODDI fitting results for the 1.25 mm isotropic resolution in vivo data. CSF volume fraction(fiso), intra-cellular volume fraction(fintra) and orientation dispersion index (ODI) from NODDI calculated from Single band reference images with 2 average, SENSE and ms-DAGER results are shown. Though bias exists in ODI maps, ms-DAGER can produce improved fitting results compared to SENSE results which are significantly corrupted by noise.

**Fig S6.**
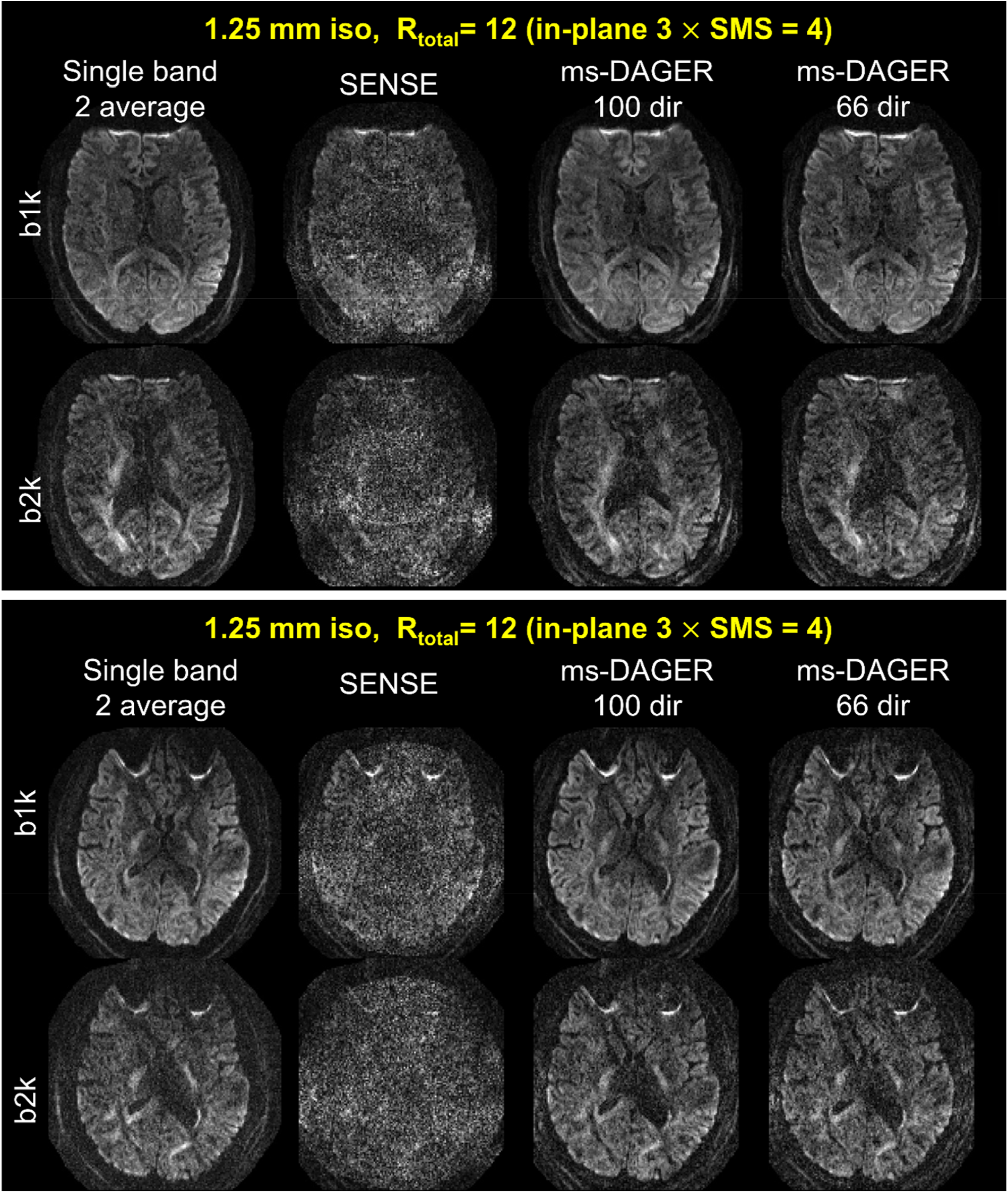
Reconstruction results for the 1.25 mm isotropic resolution in vivo data from two other subjects. Single band reference images, SENSE and ms-DAGER results are shown. b=1000s/mm^2^ (‘b1k’) and b=2000s/mm^2^ (‘b2k’) images are both shown. ms-DAGER can consistently improve image quality.

**Fig S7.**
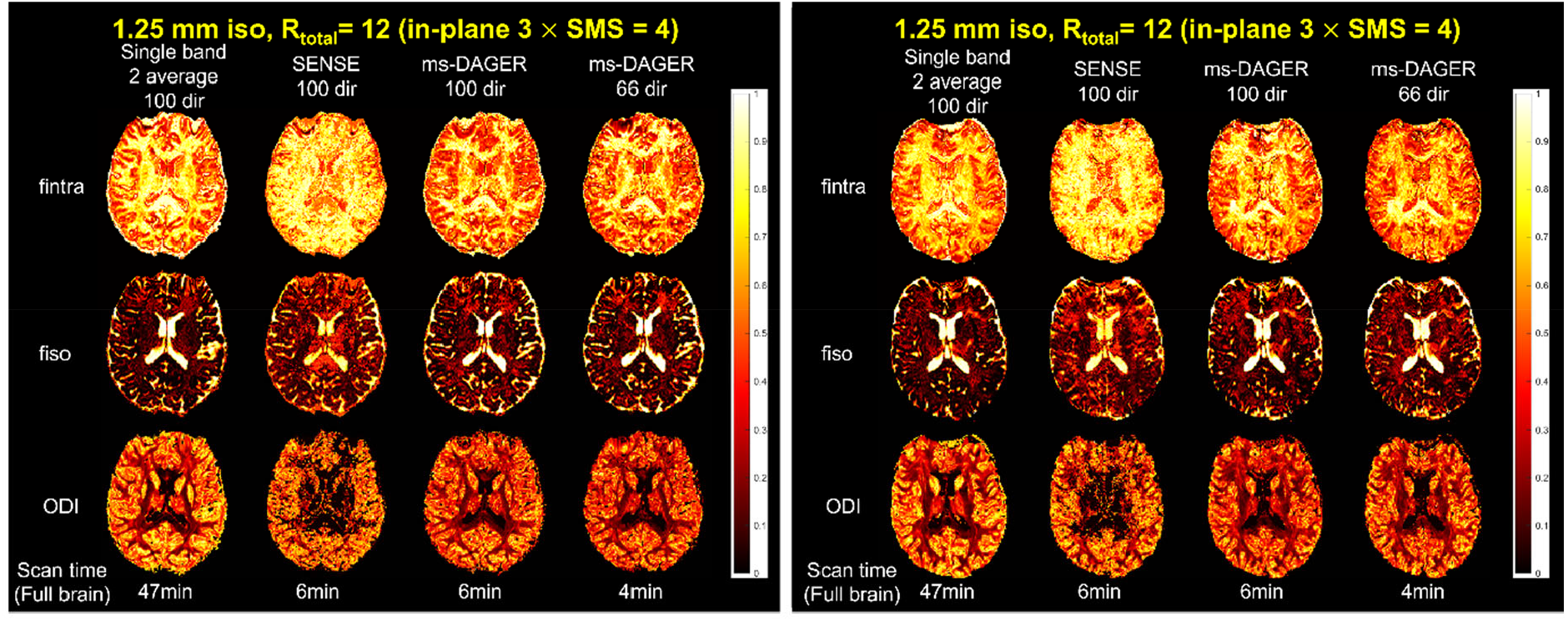
NODDI fitting results for the 1.25 mm isotropic resolution in vivo data for other subjects. CSF volume fraction(fiso), intra-cellular volume fraction(fintra) and Orientation dispersion index (ODI) from NODDI calculated from Single band reference images with 2 average, SENSE and ms-DAGER are shown are shown. Compared to SENSE, ms-DAGER consistently improve the fitting accuracy, showing high reproducibility across subjects

**Fig S8.**
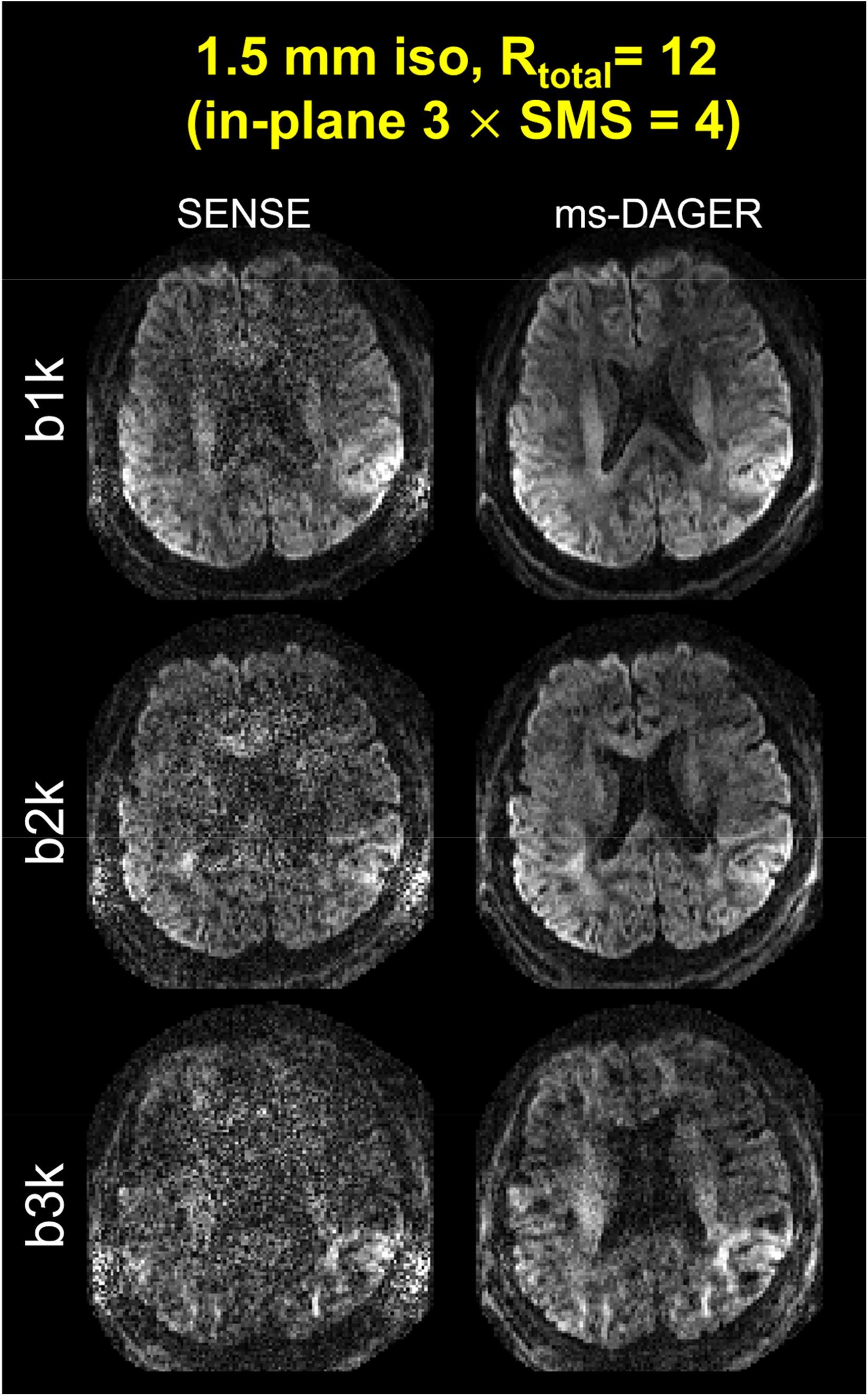
Reconstruction results for the 1.5 mm isotropic resolution 3-shell in vivo data. SENSE and ms-DAGER results are shown. b=1000s/mm^2^ (‘ b1k ‘), b=2000s/mm^2^ (‘ b2k ‘) and b=3000s/mm^2^ (‘b3k’)images are both shown. The image quality of higher b value shell can be significantly improved compared to SENSE with the help of cross-shell information sharing.

